# Does an artistic representation of brand logos increase their effect on customers?

**DOI:** 10.1101/2021.06.15.448420

**Authors:** Julia Krämer, Timo Rott, Jan-Gerd Tenberge, Patrick Schiffler, Andreas Johnen, Nils C. Landmeyer, Ferencz Olivier, Heinz Wiendl, Sven G. Meuth

## Abstract

**Background:** In numerous fMRI studies, brands strongly confound the customer’s economic decisions on a neural level by modulating cortical activity in reward-related areas.

**Objective:** To test the hypothesis that the effect of logos can be increased by artistic logo representations, we presented logos in original and artistically changed versions during fMRI.

**Methods:** Following a pre-study survey on the familiarity of original brand logos, 15 logos rated as “familiar” and 10 logos rated as “unfamiliar” were selected for fMRI experiment. During fMRI, 15 healthy subjects were presented with original and artistically changed logos out of the familiar/unfamiliar categories. A whole-brain and ROI analysis for reward-related areas were performed. Moreover, logo-induced valence and arousal were measured with the self-assessment manikin.

**Results:** Whole-brain analysis revealed activation in bilateral visual cortex for artistically changed logos (familiar/unfamiliar) compared to original logos. No significant effect could be detected for the ROI analysis. On average, the logos caused neutral emotions. However, when analyzing valence and arousal for familiar/unfamiliar and original/artistically changed logos separately, familiar original logos evoked stronger positive emotions than familiar artistically changed logos. Artistically changed logos (familiar/unfamiliar) excited participants significantly more than original logos.

**Conclusion:** Artistically changed logos elicit activation in the bilateral visual cortex but not in reward-related areas.

## 1 Introduction

Brand logos are not a modern invention. Precursors of today’s logos are the arms and heraldic symbols of ancient Greeks, Pharaohs, and Romans. Unique, easy to recognize, memorable, clear, and trustworthy brand logos help attract potential customers and clients, as strong brands “make connections” ^1^ and “promise certain advantages of a product” ^2^. Numerous functional magnetic resonance imaging (fMRI) studies in the field of consumer neuroscience, a subfield of neuroeconomics, provided evidence that customers’ brand decisions have a neural basis ^2-8^. Brands impact on “buying behavior” by specifically modulating cortical activity in the ventromedial prefrontal cortex (VMPFC), medial prefrontal cortex (MPFC), and associated limbic system ^3^. For instance, envisioning driving a culturally familiar car resulted in MPFC activity ^2^. Viewing emblems of prestigious sports and luxury brands, such as Porsche, activated the MPFC and precuneus ^9^. Erk and colleagues reported increased activation of reward-related brain areas (ventral striatum, orbitofrontal cortex, bilateral dorsolateral prefrontal cortex, anterior cingulate, and occipital regions) during the presentation of sports cars rated highest in attractiveness compared to limousines or small cars ^5^. Paulus and Frank found activity in VMPFC, anterior cingulate, and insula when participants had to choose their preferred beverage out of two familiar soft drinks ^8^. Likewise, and in analogy to the “Coca-Cola” test ^10^, McClure and colleagues described a consistent neural response in the VMPFC that correlated with subjects’ behavioral preferences for two culturally familiar soft drinks (Coca-Cola® and Pepsi®) ^7^. Accordingly, patients with cortical damage specifically involving the VMPFC did not change their normal preference bias in a taste test when exposed to brand information: Patients and neurologically healthy adults both preferred Pepsi in a blind taste test (Coca-Cola® versus Pepsi®). However, when the taste test featured brand information, patients with VMPFC damage maintained their Pepsi preference, while neurologically healthy adults suddenly preferred Coca-Cola, showing the ‘Pepsi paradox’^11^.

For sensorily nearly indistinguishable products of equal quality (coffee or beer brands), Deppe and colleagues revealed a non-linear winner-takes-all effect for a participant’s favorite brand during a binary buying decision task ^4^. This behavior was functionally characterized by increased activation in the VMPFC and other areas involved in storing, processing, and integrating self-reflection and emotions experienced during decision making – a pattern of activation only visible while choosing a favorite brand ^4^. In a binary credibility judgment task (‘true’ or ‘false’) on news magazine headlines confounded by formally decision irrelevant framing information (magazine logos), individual activity changes in the VMPFC during judgments correlated with a participant’s degree of susceptibility to framing information ^3^. In line with the somatic marker hypothesis of Damasio and colleagues, a participant’s favorite brand (be it a favorite sports car, beverage, or news magazine) evokes a somatic bioregulatory state that either “forces attention on the negative outcome of the decision” and immediately rejects the negative course of action of not choosing the first-choice brand, or, if the marker is positive, becomes a “beacon of incentive” to select the first-choice brand ^3,12^. Brands can strongly confound the *neural* basis of a customer’s economic behavior ^3^. It remains unclear, however, if this effect can be further expanded by an artistic representation of a brand logo. A higher information value through a more complex artistic logo representation – providing increased detail, a more prominent surface structure, and enhanced haptics – may trigger a stronger emotional response and, as a consequence, increase the economic value of a logo. Against this background, the present study aims to investigate whether differences in the graphic representation of logos elicit detectable differences in brain activity.

## 2 Materials and methods

### 2.1 Demographic data

15 right-handed subjects (8 female, 7 male; mean age 26 years (y), range 22–30 y; mean level of education 16.57 y) without any history of neurological, psychiatric, or internal diseases were recruited via advertising from within the Westphalian Wilhelms-University of Münster, Germany. Standard exclusion criteria for magnetic resonance imaging (MRI) examinations, such as metal implants, were applied. Due to the visual nature of the stimuli, subjects with strong myopia or other relevant visual constraints were excluded. Further exclusion criteria were: age under 18 y and over 69 y; any pre-existing medical conditions known to be associated with brain pathology; pregnancy; previous or current addiction to substances; claustrophobia; reduced general condition; familiarity with the artistic logos used in the study. We determined participants’ handedness in advance using a standard questionnaire ^24^.

### 2.2 Artistically changed logos

The artistically changed logos used in this study were selected from the artwork of a Münster based artist (see **Figure 1** for an example). Their information content was more complex than that of their original counterparts: graphic detail was increased by changing and adding to the logo’s surroundings; a relief-like surface structure was created by integrating different materials, photographs, and newspaper clippings in the artwork as pieces of a collage; further, haptics was enhanced by applying color variations and a “used” look to the image. To define the information content of artistically changed logos, we made use of the known correlation between image complexity and digital file size ^25-28^. The original and artistically changed logos were scaled to a resolution of 800 × 800 pixels. As shown in **Supplementary Figure 1**, the file size of original and artistically changed logos was significantly different (original logos: mean file size 289.64 ± 151.44 kBytes, range 74–687 kBytes; artistically changed logos: mean file size 711.88 ± 183.01, range 279–1100 kBytes; two-sided t-test, p=1.57 e^-12^).

**Figure 1:**
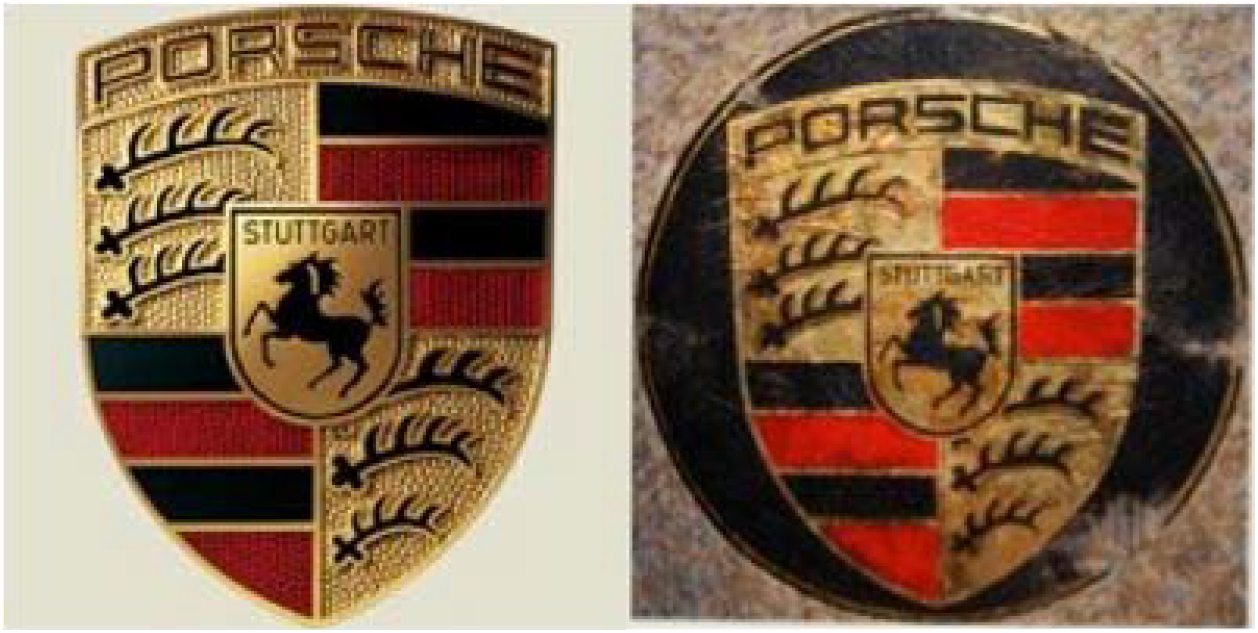
Example of an original and an artistically changed Porsche logo. Left: original brand logo, right: artistically changed logo.

### 2.3 Pre-study familiarity survey on original logos

Since the familiarity of logos can have a strong influence on how they are perceived ^4^, a pre-study familiarity survey on original brand logos was carried out. Fifteen students of the Westphalian Wilhelms-University of Münster (8 female, 7 male; mean age 23 y), other than the participants included in the study, took part in the survey. They had to judge 25 original logos as “familiar”, “unfamiliar”, or “previously encountered” (defined as *visually encountered but unaware of the affiliated brand*) (**Supplementary Figure 2**). **Figure 2** depicts the results of the survey: nine logos were rated familiar by all students, six logos were rated familiar by the majority of students, and 10 logos were rated unfamiliar by nearly all students. Based on this outcome, we categorized 15 logos as “familiar” and 10 logos as “unfamiliar” for our subsequent fMRI experiment.

**Figure 2.**
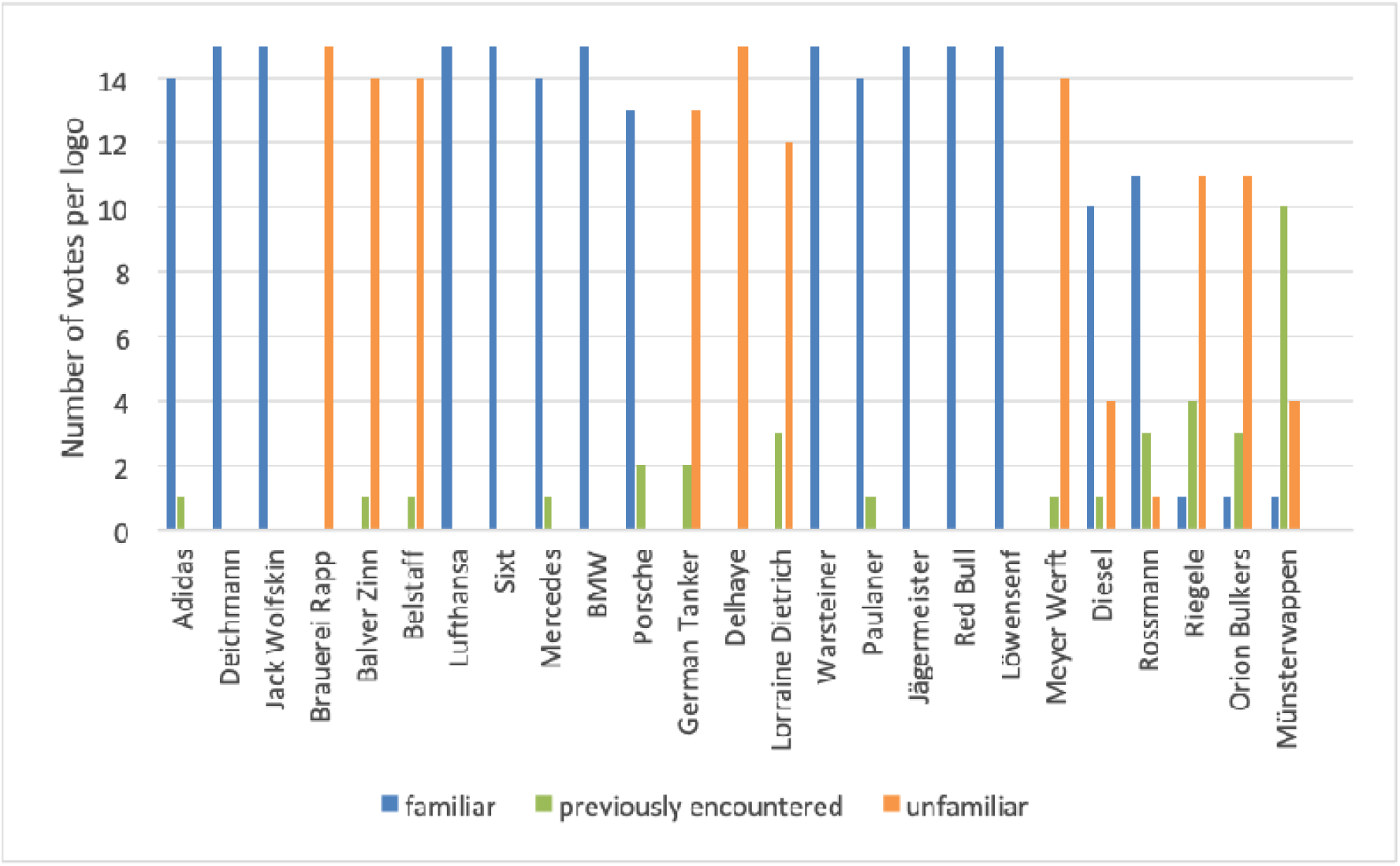
germeist wensenf were rated as “familiar” (blue) by all students (15 votes per logo). The logos Adidas, Mercedes, and Paulaner (14 votes per logo), Porsche (13 votes), Rossmann (11 votes), and Diesel (10 votes) were rated as “familiar” by the majority of students. The logos Brauerei Rapp, Delhaye, Balver Zinn, Belstaff, Meyer Werft (14 votes per logo), German Tanker (13 votes), Lorraine Dietrich (12 votes), Riegele, and Orion Bulkers (11 votes per logo) were rated as “unfamiliar” (orange) by the majority of students. The Muensterwappen was rated as “familiar” by one student and as “unfamiliar” by four students, while 10 participants stated that they had “previously encountered” (green) this logo, leading us to categorize the logo as unfamiliar.

### 2.4 Paradigm

A blocked-design was created with the neuropsychological software PsychoPy v1.84.4 (http://psychopy.org/installation.html). In the first 13 seconds (sec) of the paradigm, a short instruction explained the answer modes for the questions: *“Hello, in the following presentation you will see logos and once in a while you need to answer questions with two response possibilities: for answer R: press the right button; for answer L: press the left button”*. To answer the questions, participants were supplied with MRI compatible response boxes in their right and left hand. During fMRI, 15 experimental blocks were presented. Before the start of a new block, a fixation cross was displayed for two sec, ensuring participants would focus on the screen. Each block consisted of five logos selected from one exclusive condition (familiar-original, familiar-artwork, unfamiliar-original, unfamiliar-artwork) and presented for 5 sec each (**Figure 3**), resulting in a total length of 25 sec per block. Following the blocks 1 to 14, a two-response question referring to the last presented logo was displayed for 8 sec (**Figure 3** and **Supplementary Table 1**).

**Figure 3.**
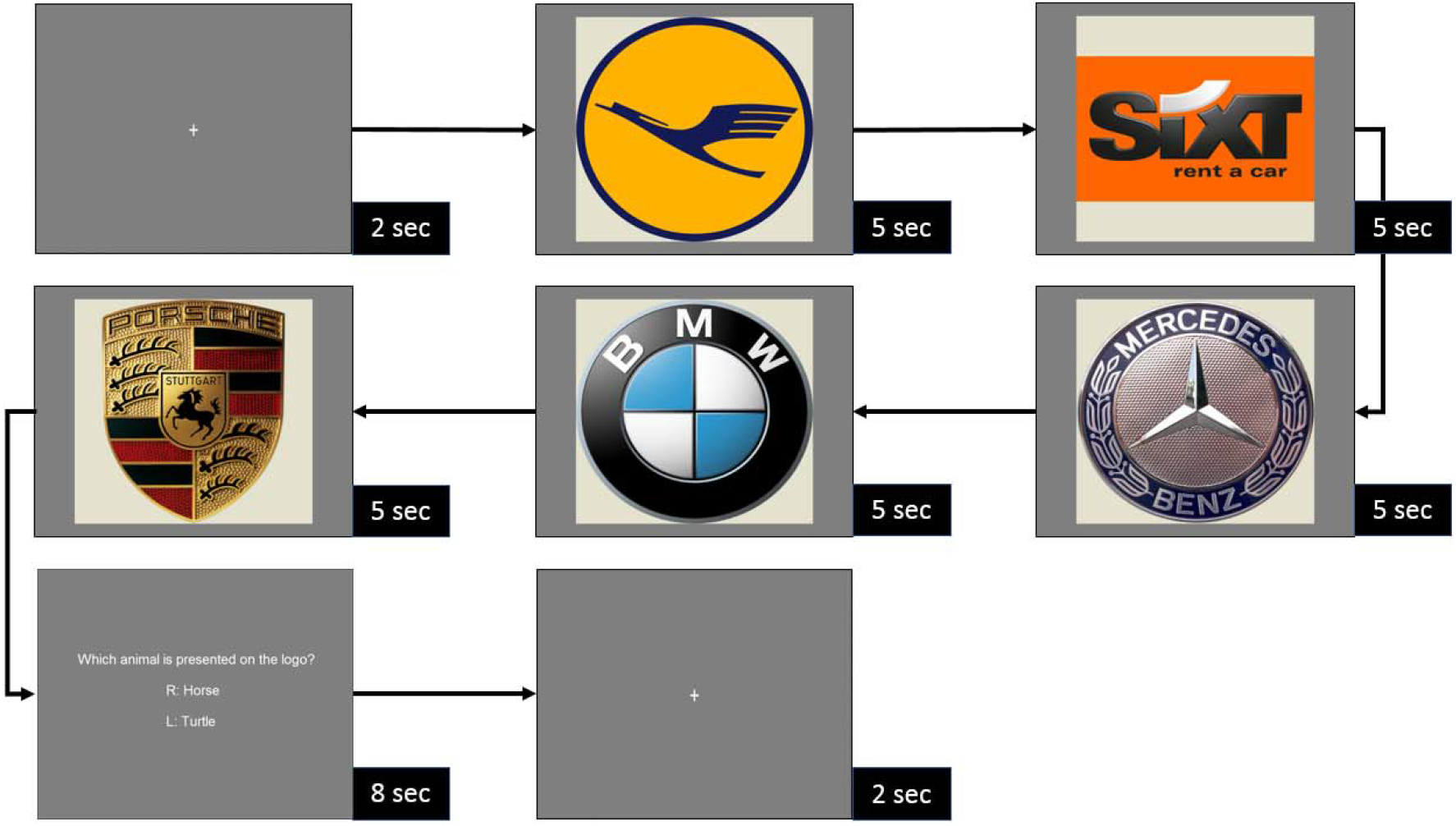
Example block of the study paradigm: After shortly displaying the fixation cross, five familiar-original logos are presented for 5 sec each. Following the block, a two-response question on the last presented logo is displayed for 8 sec, and answer possibilities are marked with “R” and “L”.

The questions were easy to answer and had the pure purpose of maintaining the participants’ attention. Participants had to answer the presented questions (forced-choice) by pressing a button on the corresponding MRI compatible response box in their right or left hand. The responses were recorded with the stimulation software PsychoPy. After block 15, no further question was presented, and the participants were informed that the first part of the study was finished.

Blocks with original brand logos and artistically changed logos were presented in turn. In the first 10 blocks, individual logos were not repeated. In blocks 11 to 15, individual logos were repeated but in a different order and different mix. **Supplementary Table 2** shows the conditions for the 15 blocks with the exact order of the respective logos.

### 2.5 Image presentation

A dedicated fMRI projection system (Covilex, Magdeburg, Germany; based on the beamer DLA-RS66e, JVC Kenwood Europe, Friedberg, Germany) provided high-quality images, guided through a large diameter RF blocking duct onto an approximately 30 × 25 cm field on a screen fixed at the rear opening of the MR bore. Lying in the scanner, participants could view the screen via a 45° mirror fixed at the top of the head coil. Triggered by the scanner, all images were presented with the neuropsychological stimulation software PsychoPy v1.84.4. To prevent confounding visual stimulation, we took care to present all logos in equal size, position, background, and luminance. Head fixation was achieved with foam pads and a soft headband. Earplugs and headsets were used to protect participants against scanner noise and to permit communication.

### 2.6 MR image acquisition

All data were acquired on a 3 T MAGNETOM Prisma^fit^ MRI scanner (Siemens AG, Erlangen, Germany), nominal gradient strength 80 mT/m, maximal slew rate 200 T/m/s. For resonance signal acquisition, the standard 20 channel head coil was used. Following a survey, a 3D isotropic T1-weighted (T1w) dataset of the whole head with a measured voxel size of 1.0 mm edge length was acquired for anatomical identification and coregistration to Talairach space, using a 3D MP-RAGE sequence in sagittal slice orientation, FOV 256 × 256 × 192 mm (frequency × phase × slice encoding in fh/ap/lr direction), acquired matrix 256 × 256 × 96, reconstructed to 192 slices. Contrast was defined by repetition time (TR) = 2130 ms, echo time (TE) = 2.28 ms, flip angle (FA) = 8°, magnetization preparation by an inversion recovery prepulse with inversion time (TI) = 900 ms, parallel imaging acceleration factor 2, echo train length 208, acquisition bandwidth (BW) per pixel 200 Hz, total acquisition time 4:56 minutes. For functional images, blood oxygenation level dependent (BOLD) contrast images were acquired using a T2*-weighted single-shot gradient echo-planar imaging (EPI) sequence that covered nearly the whole brain. The data set consisted of 33 transversal slices of 3.8 mm thickness, slice gap 0.38 mm, FOV 210 × 210 mm, matrix 64 × 64, phase encoding in ap direction. Slices were oriented parallel to the ac-pc-line. Contrast parameters were TR = 2000 ms, TE = 29 ms, FA = 90°, EPI-factor (echo train length) 64, selective fat suppression. Acquisition time per set of 33 slices was 2 sec, a total of 267 volumes were acquired, total acquisition time was 9 minutes.

### 2.7 Data analysis

Data analysis was performed using Statistical Parametric Mapping (SPM12; Wellcome Department of Cognitive Neurology, London, UK; https://www.fil.ion.ucl.ac.uk/spm/software/spm12/) to allow functional data sets to be entered into group analyses. To correct for head movements between the 33 slices, all EPI volumes were realigned to the first volume acquired using a 6-parameter affine rigid-body transformation. The calculated mean volume was saved for coregistration with the participant’s T1w structural scan. The EPI volumes were spatially normalized, warped into the Montreal Neurological Institute (MNI) EPI standard template of 152 averaged brains, and resampled to 2 × 2 × 2 mm^3^ resolution ^29^. All normalized functional volumes were smoothed with an isotropic 8-mm FWHM Gaussian kernel to decrease spatial noise. After realignment, co-registration, stereotaxic normalization, and smoothing, statistical parametric maps were calculated separately for each subject according to the hemodynamic response function. Global changes in fMRI response from scan to scan were removed by proportional scaling to have a common global mean voxel value. To correct for long-term drift effects, we applied high-pass filtering with a cut-off frequency of 0.008 Hz. Grand mean scaling was session (participant) specific. The hemodynamic responses without temporal derivatives were modeled into a block-related statistical design based on the General Linear Model (GLM) ^30^. Statistical parametric maps were calculated independently for each subject by using a boxcar regressor convolved with a hemodynamic response function (HRF). Individual, regionally specific effects of the conditions for each subject were compared using linear contrasts, resulting in t-statistics for every voxel. All contrasts calculated in the first and second level analysis are shown in **Table 1**.

**Table 1.** All contrasts calculated in the first and second level analysis.

| Contrasts of the first and second level analysis |
| --- |
| artistically changed logos > original logos |
| original logos > artistically changed logos |
| familiar: artistically changed logos > original logos |
| familiar: original logos > artistically changed logos |
| unfamiliar: artistically changed logos > original logos |
| unfamiliar: original logos > artistically changed logos |
| familiar logos > unfamiliar logos |
| unfamiliar logos > familiar logos |
| interaction: artistically changed/original logos > familiar/unfamiliar |
| interaction: familiar/unfamiliar > artistically changed/original logos |
| logos > file size |
| file size > logos |

We carried out a group analysis on a second level using a whole-brain random-effects model (one-sample t-test). The appropriate individual statistical contrast images from single-subject analyses were used for this group analysis. To account for gender differences, the participant’s gender was integrated into the analysis as a nuisance variable. In the whole-brain analysis, only those clusters that contained a minimum of five contiguous voxels with a threshold of p < 0.05 (corrected for multiple comparisons using the family-wise error (FWE) correction) were considered to reflect significant neural activation. Anatomical regions were identified using the Talairach Daemon ^31^ and the nomenclature of Brodmann ^32^.

A region of interest (ROI) analysis was performed using the SPM-Toolbox WFU-Pickatlas (Version 3.0.5, https://www.fil.ion.ucl.ac.uk/spm/ext/#WFU_PickAtlas) ^33,34^. With this method, ROI masks based on the Talairach Daemon database were generated for the nucleus accumbens, the ventral tegmental area, the ventral striatum, the ventral pallidum, the prefrontal cortex, the amygdala, the hippocampus, and the thalamus. Only those clusters with a size > 6 voxels and p < 0.001 (corrected for multiple comparisons using the FWE correction) were considered significant in the ROI analysis ^35-38^.

### 2.8 Self-assessment manikin and survey on the familiarity of the original logos

After the fMRI experiment, participants’ valence and arousal to all original and artistically changed logos were measured with the self-assessment manikin (SAM) ^39^. The SAM is a brief, nonverbal, culture-independent, and picture-oriented questionnaire developed to measure the features valence, arousal, and dominance associated with a person’s affective reaction to a wide variety of stimuli ^40,41^. Both dimensions (valence and arousal) are represented by five pictograms, respectively. Participants had to decide between nine fields for each emotion (fields 1–2= high level of valence/arousal; fields 3–6= neutral level of valence/arousal; fields 7–10= no or low level of valence/arousal), marking the selected field with a cross (**Supplementary Figure 3**). Moreover, the subjects were requested to judge the familiarity of the original logos on a five-point Likert scale in graphic form (**Supplementary Figure 4**).

### 2.9 Statistical analyses

Statistical analysis of behavioral data was performed using Microsoft Excel 2016 (Microsoft Corp., Redmond, USA). Differences in SAM ratings for valence and arousal between original and artistically changed logos, between familiar original and familiar artistically changed logos, and between unfamiliar original and unfamiliar artistically changed logos were analyzed using two-sided t-tests. Differences in file size between original brand logos and artistically changed logos were tested using two-sided t-tests. Differences were considered statistically significant if p ≤ 0.05.

## 3 Results

### 3.1 Results of the whole-brain analysis

Compared to original brand logos, artistically changed logos elicited activation in the bilateral visual cortex (Brodmann areas (BA) 18) (MNI coordinates [12, -86, -4], peak t value = 14.18, extent threshold = 382 voxels) **(Figure 4A)**. The contrast original logos > artistically changed logos revealed no significant differences in activation patterns. When analyzing familiar and unfamiliar logos separately, artistically changed logos elicited activation in the bilateral visual cortex compared to original brand logos (contrast familiar: artistically changed logos > original logos: bilateral BA 18, MNI coordinates [12, -86, -4], peak t value = 11.32, extent threshold = 120 voxels (**Figure 4B**); contrast unfamiliar: artistically changed logos > original logos: bilateral BA 18, MNI coordinates [10, -86, -4], peak t value = 14.17, extent threshold = 219 voxels (**Figure 4C**)). The contrasts familiar logos > unfamiliar logos and unfamiliar logos > familiar logos revealed activation in the visual cortex only when lowering the threshold to 0.001 (uncorrected for multiple comparisons). Compared to unfamiliar logos, familiar logos additionally elicited activation in the left BA 21 (threshold at 0.001, uncorrected for multiple comparisons). The contrast interaction: artistically changed/original logos > familiar/unfamiliar revealed activation in BA 6, 7, 10, 17 and 18 (threshold at 0.001, uncorrected for multiple comparisons). The contrast interaction: familiar/unfamiliar > artistically changed/original logos revealed no differences in activation, neither for the threshold 0.05 nor for the threshold 0.001 (uncorrected for multiple comparisons). File size, viewed in isolation, elicited significant activation in the bilateral visual cortex (BA 18) (MNI coordinates [22, -78, -10], peak t value = 21.89, extent threshold = 959 voxels) (**Figure 4D**). The contrast logos > file size revealed activation of left BA 7 (MNI coordinates [-6, -64, -54], peak t value = 10.92, extent threshold = 44 voxels) and 21 (MNI coordinates [-62, -30, 0], peak t value = 11.30, extent threshold = 92 voxels). For the contrast file size > logos, activation in the bilateral visual cortex was eliminated.

**Figure 4.**
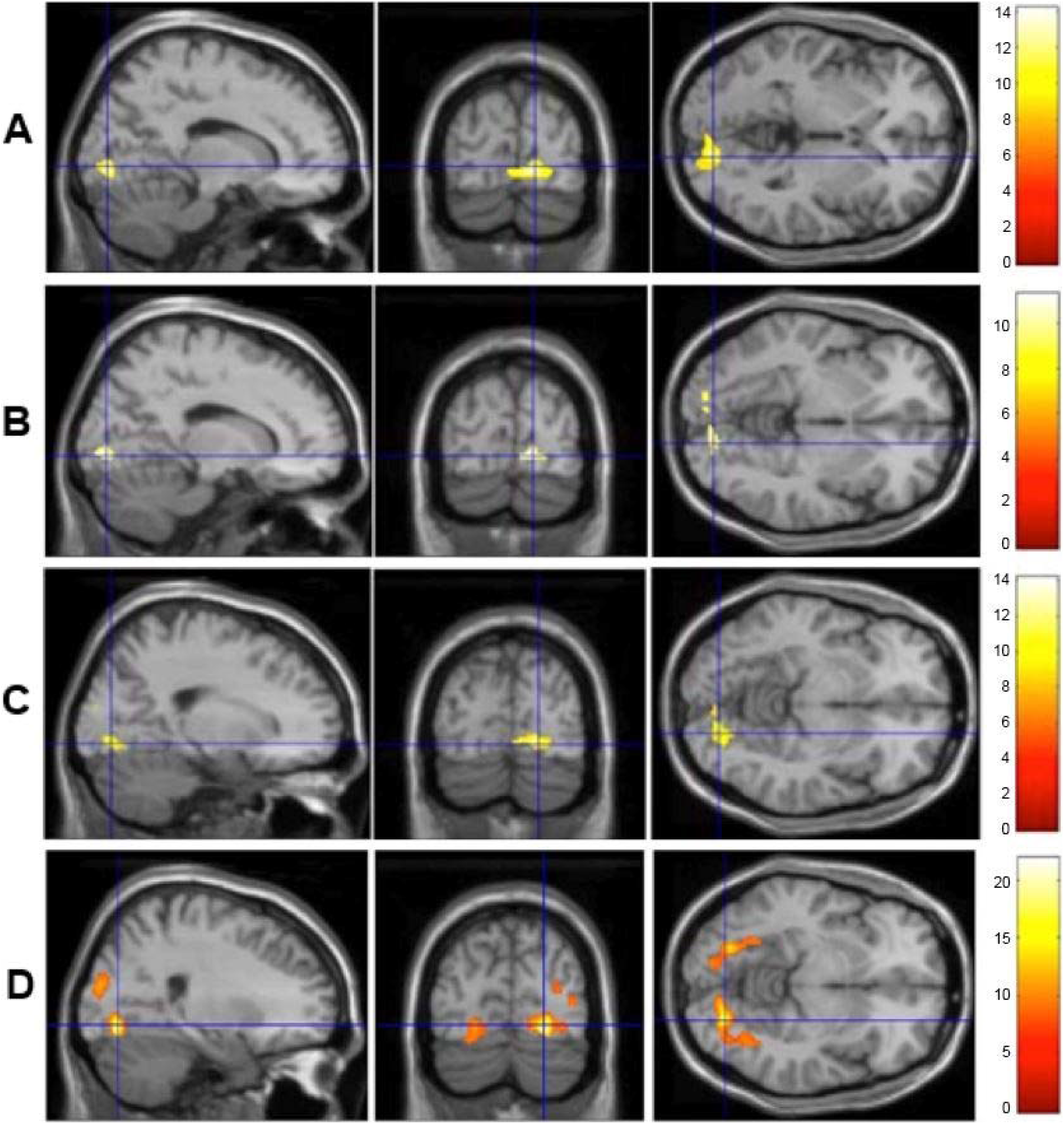
**(A)** Artistically changed logos elicited activation in the bilateral visual cortex compared to original brand logos. Areas of significant fMRI signal change for the contrast artistically changed logos > original logos are shown as color overlays on the T1-MNI reference brain (threshold = p < 0.05, FWE correction). The colored bar indicates the t statistics of the activation. **(B)-(C)** When analyzing familiar **(B)** and unfamiliar logos **(C)** separately, artistically changed logos elicited activation in the bilateral visual cortex compared to original brand logos. Areas of significant fMRI signal change for the contrasts familiar: artistically changed logos > original logos **(B)** and unfamiliar: artistically changed logos > original logos **(C)** are shown as color overlays on the T1-MNI reference brain (threshold = p < 0.05, FWE correction). The colored bar indicates the t statistics of the activation. **(D)** The logos’ file sizes elicited significant activation in the bilateral visual cortex. Areas of significant fMRI signal change for the contrast file size are shown as color overlays on the T1-MNI reference brain (threshold = p < 0.05, FWE correction). The colored bar indicates the t statistics of the activation.

### 3.2 fMRI results of the ROI analysis

While some participants demonstrated activities in areas of the reward system in the first-level analysis for several contrasts, especially for the contrast artistically changed logos > original logos, the second-level analysis revealed no significant effect for any contrast.

### 3.3 Survey on the familiarity of the original logos

Values > 3 indicate a familiar logo and values < 3 an unfamiliar logo. With results comparable to the pre-study survey on the familiarity of the original brand logos, the same 15 logos were rated as “familiar” (mean value 4.81, SD 0.43), and the same 10 logos as “unfamiliar” (mean value 1.58, SD 1.10). Among the logos rated as “familiar”, the brands Rossmann, Adidas, and Mercedes showed a large variation of rating values (**Supplementary Figure 5**).

### 3.4 Self-assessment manikin ratings

As demonstrated in **Figure 5**, the logos used, whether original or artistically changed, caused neither positive nor negative valence in the participants, and mostly neutral arousal (mean valence of original logos 5.76 ± 1.31 and of artistically changed logos 5.68 ± 1.30; mean arousal of original logos 4.11 ± 2.06 and of artistically changed logos 4.58 ± 2.12). However, original logos generally elicited more positive valence in participants than artistically changed logos, though without reaching significance (p=0.065), and artistically changed logos excited subjects significantly more than original logos (p=0.0065) (**Figure 5**).

**Figure 5:**
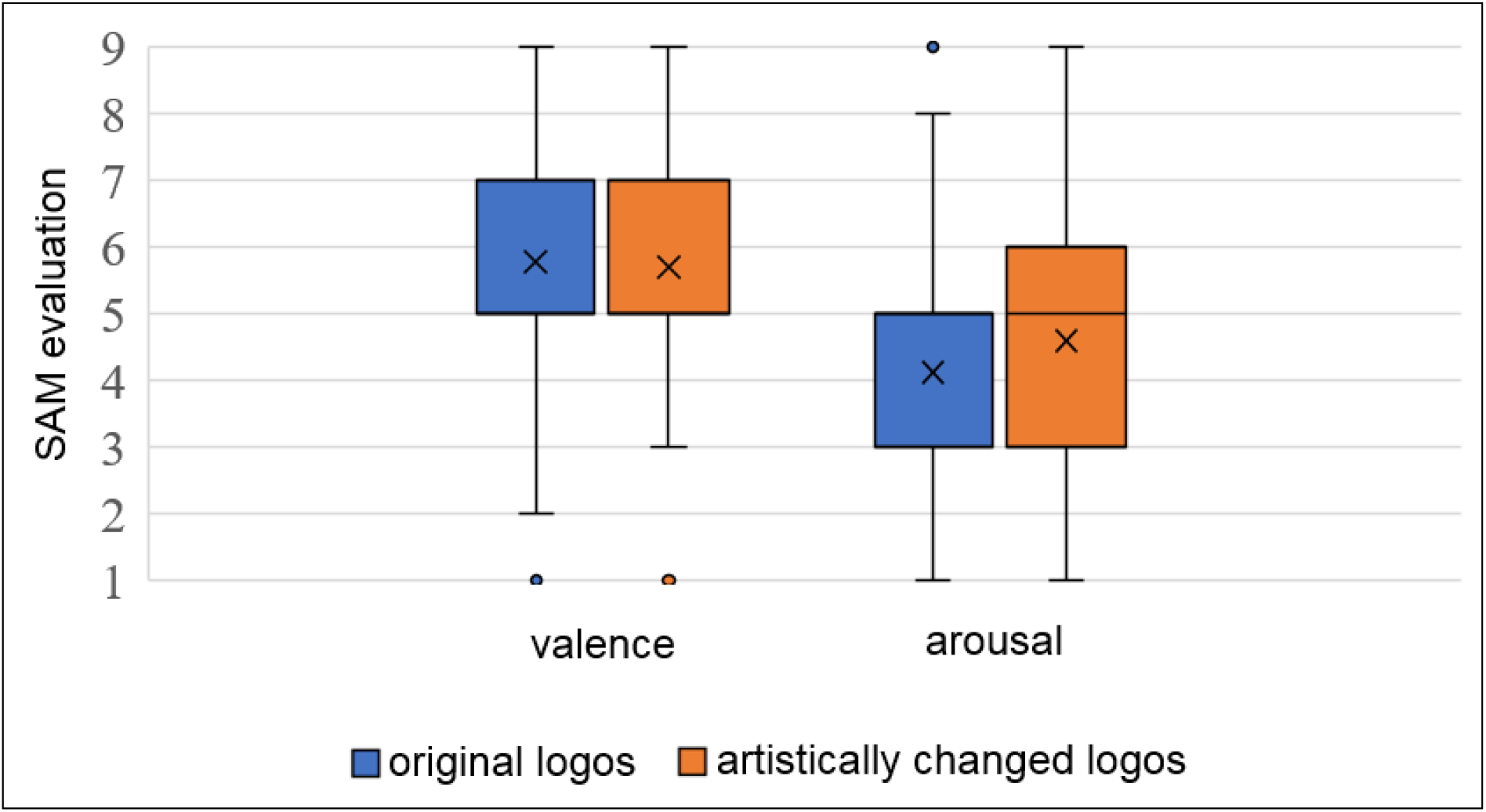
Boxplots depicting results of the SAM evaluation for valence and arousal for original (blue) and artistically changed logos (orange). × presents the mean value, ° presents outliers. 50% of all evaluations lie within the box; the median value is 5.

When analyzing the induced valence/arousal for familiar/unfamiliar and original/artistically changed logos separately, familiar original logos evoked more positive valence than familiar artistically changed logos (p=0.006) (**Table 2A**). Unfamiliar original and unfamiliar artistically changed logos caused almost identical neutral valence (p=0.269) (**Table 2A**). Familiar artistically changed logos excited subjects significantly more than familiar original logos (p=0.007), and unfamiliar artistically changed logos excited subjects significantly more than unfamiliar original logos (p=0.0001) (**Table 2B**).

**Table 2A.**
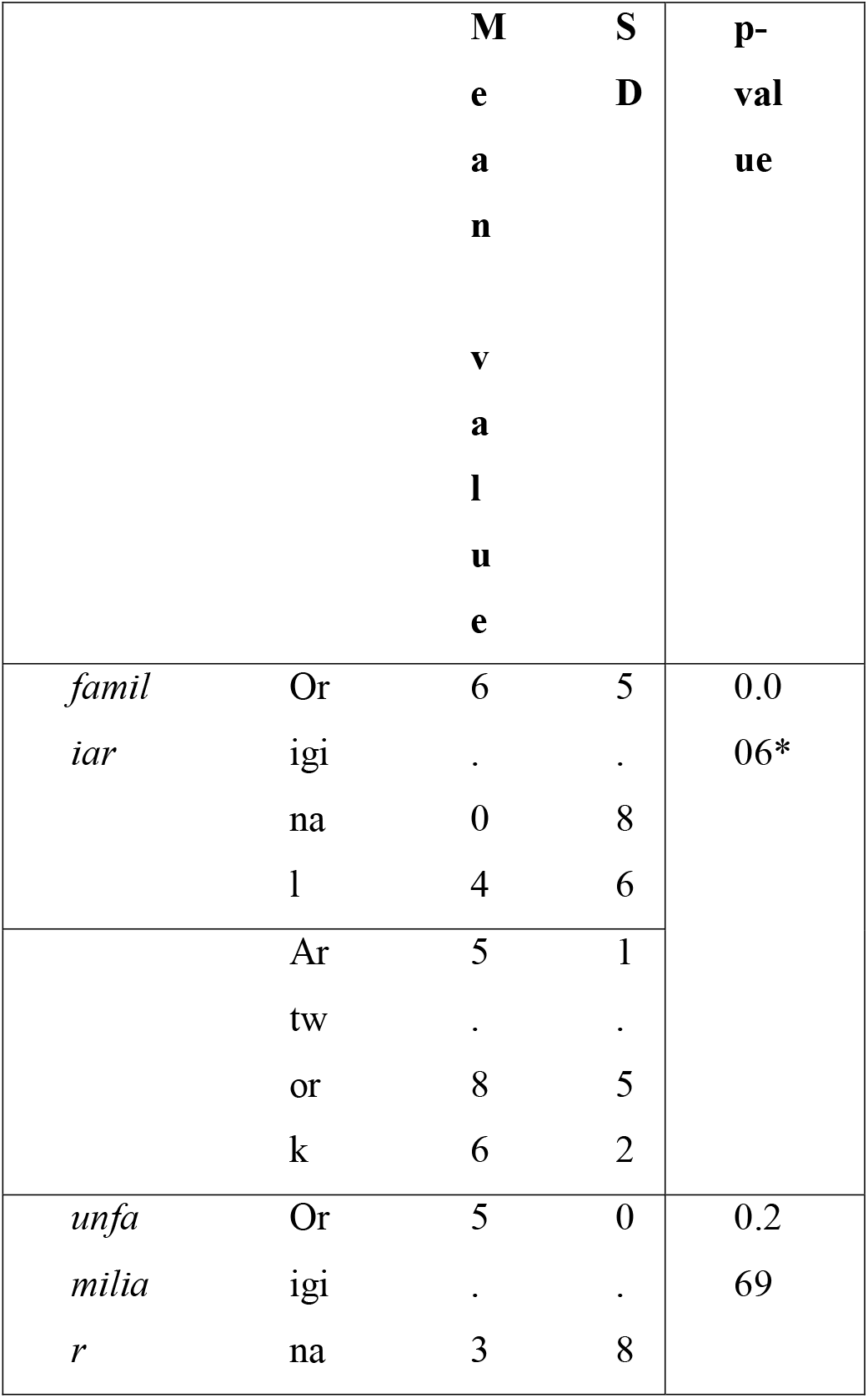
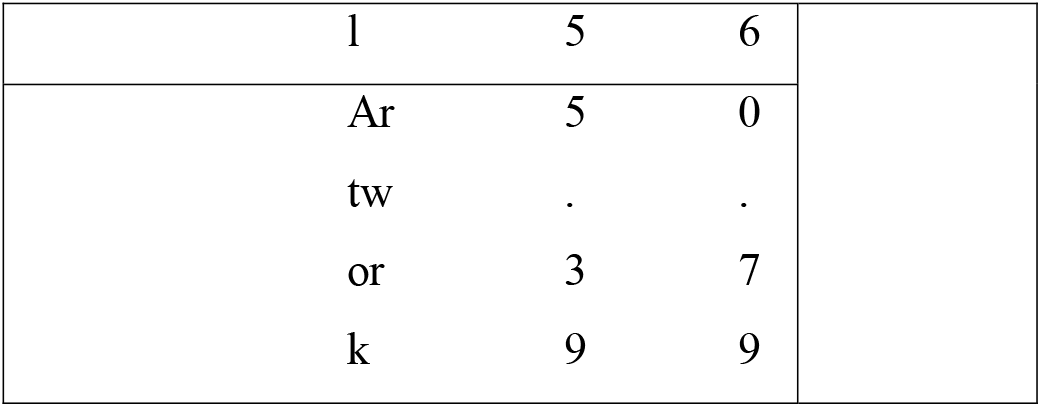
Results of the t-test for the SAM evaluation of valence for familiar/unfamiliar and original/artistically changed logos.

|  |  | **M** | **S** | **p-** |
| --- | --- | --- | --- | --- |
|  |  | **e** | **D** | **val** |
|  |  | **a** |  | **ue** |
|  |  | **n** |  |  |
|  |  | **v** |  |  |
|  |  | **a** |  |  |
|  |  | **l** |  |  |
|  |  | **u** |  |  |
|  |  | **e** |  |  |
| *famil*<br>*iar* | Or | 6 | 5 | 0.0<br>06* |
|  | igi | . | . |  |
|  | na | 0 | 8 |  |
|  | l | 4 | 6 |  |
|  | Ar | 5 | 1 |  |
|  | tw | . | . |  |
|  | or | 8 | 5 |  |
|  | k | 6 | 2 |  |
| *unfa*<br>*milia*<br>*r* | Or | 5 | 0 | 0.2<br>69 |
|  | igi | . | . |  |
|  | na | 3 | 8 |  |

|  |  |  |
| --- | --- | --- |
| l | 5 | 6 |
| Ar | 5 | 0 |
| tw | . | . |
| or | 3 | 7 |
| k | 9 | 9 |

**Table 2B.** Results of the t-test for the SAM evaluation of arousal for familiar/unfamiliar and original/artistically changed logos.

|  |  | Mean<br>value | SD | p-value |
| --- | --- | --- | --- | --- |
| *familiar* | Original | 4.33 | 2.19 | 0.007* |
|  | Artwork | 4.84 | 2.21 |  |
| *unfamiliar* | Original | 3.79 | 1.79 | 0.0001* |
|  | Artwork | 4.19 | 1.9 |  |

## 4 Discussion

By presenting original logos and artistically changed logos of familiar and unfamiliar brands during fMRI and measuring participants’ valence and arousal to the same logos using SAM ratings, we could demonstrate that artistically changed logos provoked an activation increase in the secondary visual cortex (BA 18) compared to original brand logos. The BA 18 play an important role in the specific structure, shape, and pattern analysis of visual stimuli ^13^. The increased cortical activity in BA 18 could be explained by the higher image complexity of the artistically changed logos (**Supplementary Figure 1**), providing more detail, surface structure, and increased haptics. This explanation is supported by the file-size dependent activation increase in the bilateral visual cortex (BA 18). The stronger excitement caused by artistically changed logos compared to original logos, whether familiar or unfamiliar, (p=0.0065) also supports the activation patterns in the bilateral visual cortex, as numerous recent studies could demonstrate that emotionally loaded visual stimuli lead to increased activation in the visual cortex ^14-16^. Because both original logos and artistically changed logos provoked neutral emotions on average (mean values of 3–6 in the SAM), we can assume that the majority of our participants were not strongly affected by our choice of logos. This would fit the lack of activation in reward-related brain areas while viewing the logos. In a few participants, however, the presented logos provoked strong valence and arousal (**Figure 5**), in the same way as for some participants, artistically changed logos elicited cortical activation in reward-related brain areas in the first-level analysis. Perhaps our fMRI paradigm, in which subjects only watched but did not evaluate or decide between different original and artistically changed logos of familiar and unfamiliar brands, was not suitable to trigger specific emotional associations based on the presented logos. In fMRI studies that revealed cortical activation in the VMPFC, MPFC, and associated limbic system, participants were requested to imagine driving a car while logos of car companies were presented ^2^, or to buy one of two presented beverages of the same product type but different brand ^4^, or to judge the credibility of newspaper headlines of different magazine brands ^3^, or to rate the attractiveness of presented cars ^5^. We decided to perform the SAM rating *after* the fMRI experiment as we intended participants to consider the presented logos unbiased and value-neutral without knowing the artistic style of the changed logos in advance. Other studies have argued that only participants’ preferred brands ^4,7,8^ and emotionally loaded visual stimuli ^17-21^ can evoke activity in reward-related brain regions. As shown by the SAM evaluations, our stimuli were not emotionally loaded (**Figure 5**). Perhaps preselecting participants with strong brand awareness of the presented logos ^22^ or connecting the changed logos with brand typical items such as cars, bottles, or clothes to evoke stronger emotions would have evoked reward-related activity. Moreover, the authenticity of the artwork used was perhaps lost due to the chosen presentation format, where participants viewed the screen via a 45° mirror while lying in the scanner. Thus, the surface profile of artistically changed logos might have been hard to perceive, and the coloring might not have been realistically presented. Unfortunately, there was no ideal presentation solution. Further, prior to the fMRI examination, the only context information our participants received was that they would view original logos and artistically changed logos. A recent fMRI study demonstrated that aesthetic judgments, like most judgments, depend on the context ^23^. Using the same database of artwork and only changing the labeling of images as being either sourced from an art gallery or computer-generated, Kirk and colleagues revealed that artwork sourced from a famous Danish art gallery was rated significantly better than artwork that was computer-generated ^23^. This contextual modulation correlated with activity in the medial orbitofrontal cortex and prefrontal cortex. The recruitment of prefrontal and orbitofrontal cortices during aesthetic judgments was shown to be significantly biased by subjects’ prior expectations about the likely hedonic value of stimuli according to their source ^23^. The main limitation of our study was the small number of participants. Therefore, our results have to be interpreted with caution. Due to the small number of participants, gender differences could not be analyzed, whereas the influence of participants’ gender was considered in the fMRI analysis.

## 5 Conclusion

Artistically changed logos elicit activation in the bilateral visual cortex due to their higher visual load, increased image complexity, and a higher level of arousal compared to original logos. The reward system, i.e., the prefrontal cortex and associated limbic system, however, is not involved. Further fMRI studies with an adapted study paradigm and larger sample size are needed to investigate how an artistic reappraisal of logos influences consumers’ behavior, potentially leading to an economic benefit.

## 8 Author Contributions

JK, TR, JGT, PS, AJ, HW, and SGM conceived the study and defined the concept. TR recruited the participants. JK, TR, JGT, PS performed the fMRI experiments, collected, analyzed, and interpreted the data, and performed confirmatory statistical analysis. TR prepared the figures and tables. JK wrote the initial draft of the manuscript. JK, TR, JGT, PS, NCL, AJ, FO, HW, and SGM critically discussed the data, revised the manuscript for intellectual content, and approved the version to be published. All authors agreed to be accountable for all aspects of the work in ensuring that questions related to the accuracy or integrity of any part of the work are appropriately investigated and resolved. HW and SGM supervised the study. HW acquired funding for the study.

## 10 Abbreviations

BA: Brodmann area
BOLD: blood oxygenation level dependent
BW: bandwidth
EPI: echo-planar imaging
FA: flip angle
fMRI: functional magnetic resonance imaging
FOV: field of view
FWE: family-wise error
GLM: general linear model
HRF: hemodynamic response function
MNI: Montreal Neurological Institute
MPFC: medial prefrontal cortex
MRI: magnetic resonance imaging
ROI: region of interest
SAM: self-assessment manikin
sec: seconds
SPM: statistical parametric mapping
TE: echo time
TI: inversion time
TR: repetition time
T1w: T1-weighted
VMPFC: ventromedial prefrontal cortex
y: years.

## 11 Acknowledgments

We thank all subjects for participating in this study. We thank Dr. Zoë Hunter of the Institute of Translational Neurology, University Hospital Münster, for proofreading the manuscript, Ima Trempler of the Institute of Psychology, Westphalian Wilhelms-University of Münster, and Dr. rer. medic Jochen Bauer of the Translational Research Imaging Center, Westphalian Wilhelms-University of Münster, for supporting performance of the fMRI experiments.

## 13 Availability statement

The datasets generated during and/or analysed during the current study are available from the corresponding author on reasonable request.

## 14 Ethical approval

The study was approved by the interdisciplinary ethics committee of the University of Münster and the Physicians’ Chamber of Westphalia-Lippe (Ärztekammer Westfalen-Lippe, 2015-155-f-S). All procedures performed in this study were in accordance with the 1964 Helsinki declaration and its later amendments or comparable ethical standards.

## 15 Consent for publication

Patients were informed of the study content in both oral and written form. All subjects gave written informed consent before participating in this study.

