## Supplemental Tables and Figures for "Does an artistic representation of brand logos increase their effect on customers?"

Supplementary Material

### 1 Supplementary Tables

| **Block** | **Logo** | **Question** |
| --- | --- | --- |
| 1 | 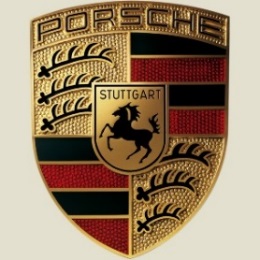 | Which animal is shown on the logo?  **R: Horse** L: Turtle |
| 2 | 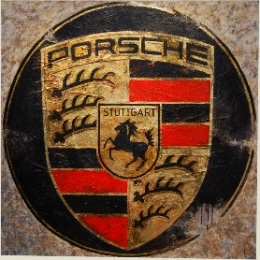 | Which color is the circle around the logo?  R: Pink **L: Black** |
| 3 | 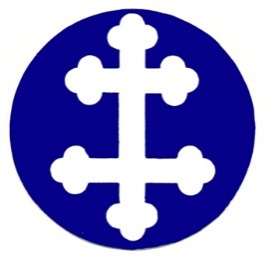 | How many colors are in this logo?  R: 4 **L: 2** |
| 4 | 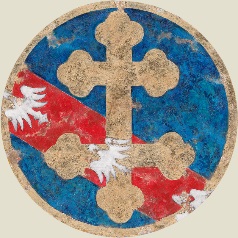 | Which animals are shown on the logo?  **R: Birds** L: Bears |
| 5 | 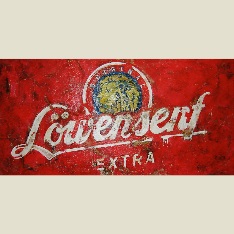 | Which color is the background?  R: Pink **L: Red** |
| 6 | 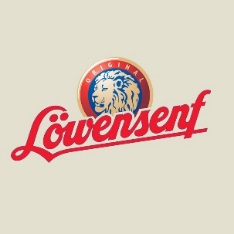 | Which color is the lion head?  **R: Golden** L: Green |
| 7 | 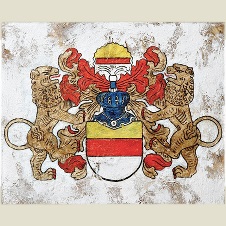 | How many lions are presented on the logo?  R: 5 **L: 2** |
| 8 | 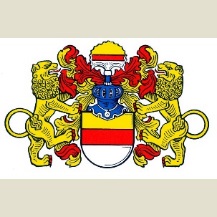 | Which color is the crown in the center of the logo?  **R: Blue** L: Grey |
| 9 | 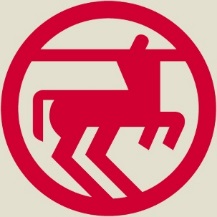 | Which hybrid creature is presented? A mixture of:  **R: Horse + human** L: Dog + cat |
| 10 | 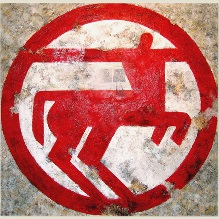 | Is the line of the circle complete or interrupted?  R: complete **L: interrupted** |
| 11 | 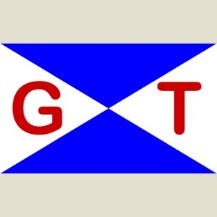 | Which letters are presented on the logo?  **R: G T** L: X Y |
| 12 | 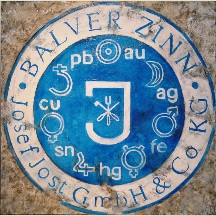 | Which letter is presented in the center of the logo?  R: Q **L: J** |
| 13 | 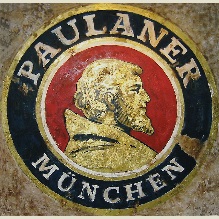 | Which color is the lettering “Paulaner München”?  R: Green **L: White** |
| 14 | 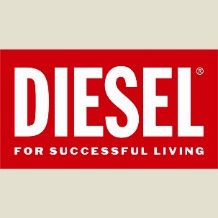 | How many colors are in this logo?  **R: 2** L: 6 |

**Supplementary Table 1** Overview of the final logo presented in each block (1–14) and the corresponding two-response question. The subjects were instructed to answer the presented question by pressing a button on the corresponding MRI compatible response box in their right (R) or left hand (L). Here, the correct answers are highlighted in **bold**.

| Block | Condition | Logo |
| --- | --- | --- |
| 1 | Familiar - Original | Lufthansa, Sixt, Mercedes, BMW, Porsche |
| 2 | Familiar – Artwork | Lufthansa, Sixt, Mercedes, BMW, Porsche |
| 3 | Unfamiliar – Original | German Tanker, Orion Bulkers, Meyer Werft, Delhaye, Lorraine Dietrich |
| 4 | Unfamiliar – Artwork | German Tanker, Orion Bulkers, Meyer Werft, Delhaye, Lorraine Dietrich |
| 5 | Familiar – Artwork | Warsteiner, Paulaner, Jägermeister, Red Bull, Löwensenf |
| 6 | Familiar – Original | Warsteiner, Paulaner, Jägermeister, Red Bull, Löwensenf |
| 7 | Unfamiliar – Artwork | Brauerei Rapp, Brauerei Riegele, Balver Zinn, Belstaff, Muensterwappen |
| 8 | Unfamiliar – Original | Brauerei Rapp, Brauerei Riegele, Balver Zinn, Belstaff, Muensterwappen |
| 9 | Familiar – Original | Adidas, Deichmann, Diesel, Jack Wolfskin, Rossmann |
| 10 | Familiar – Artwork | Adidas, Deichmann, Diesel, Jack Wolfskin, Rossmann |
| 11 | Unfamiliar – Original | Balver Zinn, Belstaff, Orion Bulkers, Brauerei Rapp, German Tanker |
| 12 | Unfamiliar – Artwork | German Tanker, Brauerei Rapp, Orion Bulkers, Belstaff, Balver Zinn |
| 13 | Familiar – Artwork | Diesel, Porsche, Lufthansa, Rossmann, Paulaner |
| 14 | Familiar – Original | Paulaner, Rossmann, Lufthansa, Porsche, Diesel |
| 15 | Unfamiliar - Artwork | Meyer Werft, Delhaye, Riegele, Lorraine Dietrich, Muensterwappen |

**Supplementary Table 2** Conditions for the 15 blocks (out of the categories: familiar-original, unfamiliar-original, familiar-artwork, unfamiliar-artwork) with their respective logos according to their sequence in the paradigm.

### 2 Supplementary Figures

File size in kBytes

|  | original logo | artistically changed logo |
| --- | --- | --- |
| Jack Wolfskin | 223 | 794 |
| Diesel | 143 | 452 |
| Balver Zinn | 422 | 757 |
| Belstaff | 157 | 784 |
| Lufthansa | 170 | 649 |
| Deichmann | 74 | 279 |
| Rossmann | 148 | 681 |
| Brauerei Rapp | 499 | 866 |
| Riegele | 442 | 772 |
| Münsterwappen | 496 | 576 |
| Sixt | 187 | 554 |
| Mercedes | 687 | 1000 |
| Porsche | 519 | 795 |
| German Tanker | 97 | 534 |
| Delhaye | 308 | 722 |
| Warsteiner | 227 | 875 |
| Jägermeister | 415 | 930 |
| Löwensenf | 223 | 453 |
| BMW | 233 | 749 |
| Orion Bulkers | 165 | 556 |
| Adidas | 270 | 595 |
| Lorraine Dietrich | 304 | 901 |
| Paulaner | 388 | 757 |
| Red Bull | 245 | 1100 |
| Meyer Werft | 199 | 666 |

**Supplementary Figure 1** File sizes (kBytes) of the 25 original logos and 25 artistically changed logos. The file sizes of artistically changed logos were significantly larger than the file sizes of original logos (original logos: mean file size 289.64 ± 151.44 kBytes, range 74–687 kBytes; artistically changed logos: mean file size 711.88 ± 183.01, range 279–1100 kBytes; two-sided t-test, p=1.57 e^-12^).

| □ familiar  □ previously encountered  □ unfamiliar | 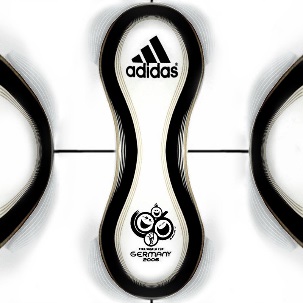 | 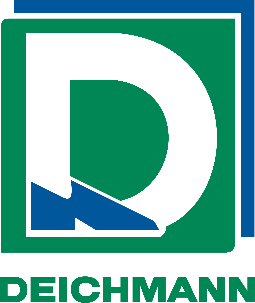 | □ familiar  □ previously encountered  □ unfamiliar |
| --- | --- | --- | --- |
| □ familiar  □ previously encountered  □ unfamiliar | 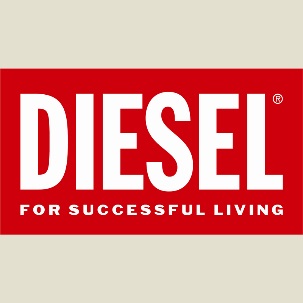 | 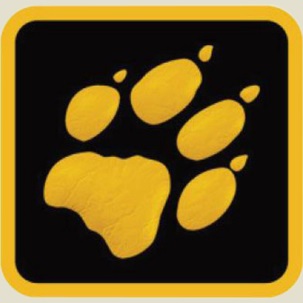 | □ familiar  □ previously encountered  □ unfamiliar |
| □ familiar  □ previously encountered  □ unfamiliar | 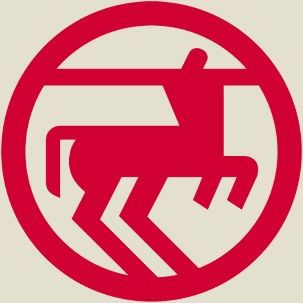 | 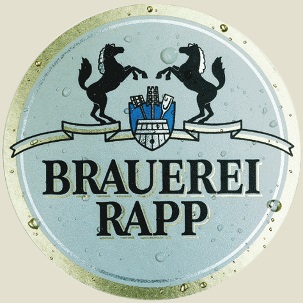 | □ familiar  □ previously encountered  □ unfamiliar |
| □ familiar  □ previously encountered  □ unfamiliar | 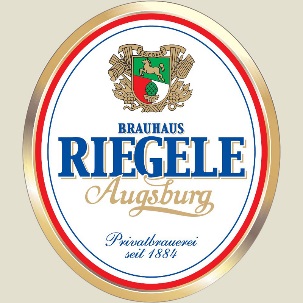 | 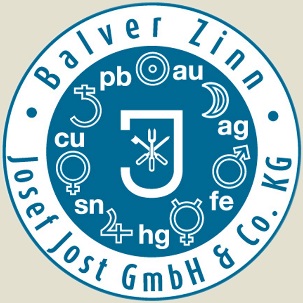 | □ familiar  □ previously encountered  □ unfamiliar |
| □ familiar  □ previously encountered  □ unfamiliar | 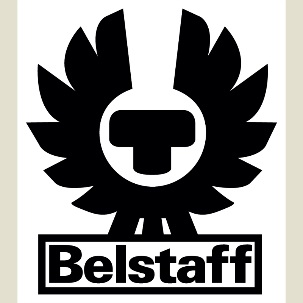 | 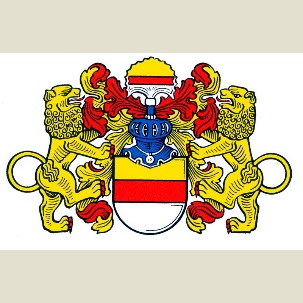 | □ familiar  □ previously encountered  □ unfamiliar |
| □ familiar  □ previously encountered  □ unfamiliar | 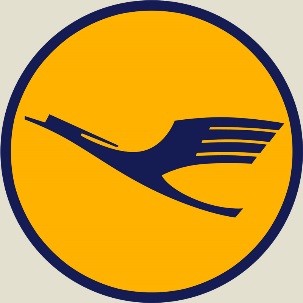 | 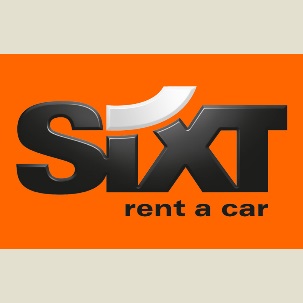 | □ familiar  □ previously encountered  □ unfamiliar |
| □ familiar  □ previously encountered  □ unfamiliar | 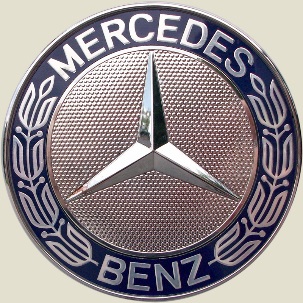 | 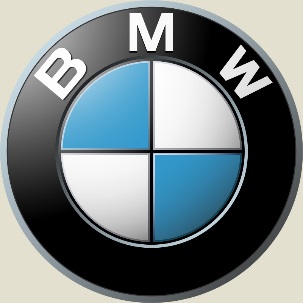 | □ familiar  □ previously encountered  □ unfamiliar |
| □ familiar  □ previously encountered  □ unfamiliar | 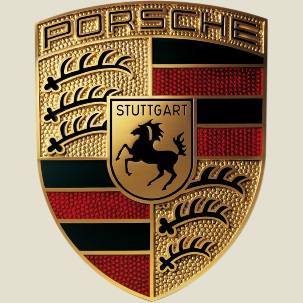 | 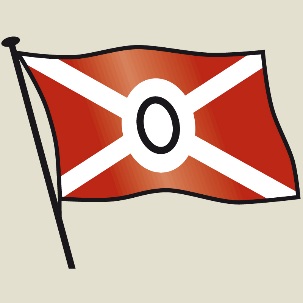 | □ familiar  □ previously encountered  □ unfamiliar |
| □ familiar  □ previously encountered  □ unfamiliar | 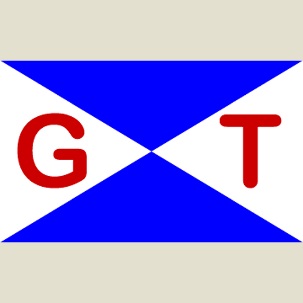 | 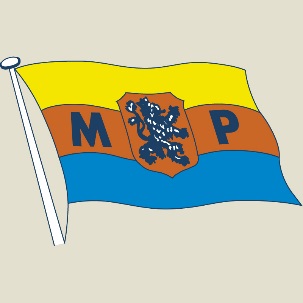 | □ familiar  □ previously encountered  □ unfamiliar |
| □ familiar  □ previously encountered  □ unfamiliar | 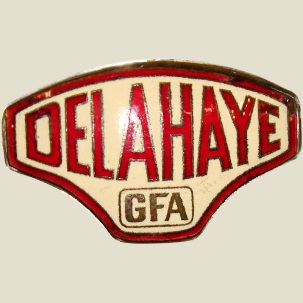 | 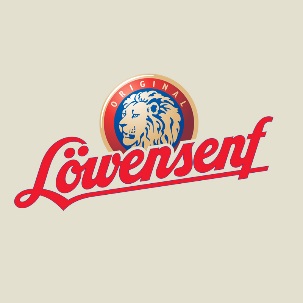 | □ familiar  □ previously encountered  □ unfamiliar |
| □ familiar  □ previously encountered  □ unfamiliar | 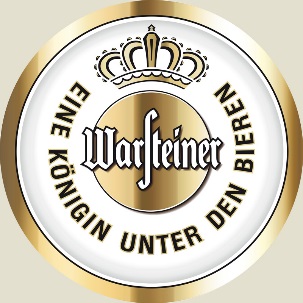 | 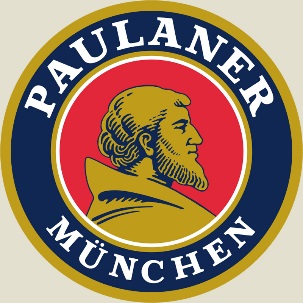 | □ familiar  □ previously encountered  □ unfamiliar |
| □ familiar  □ previously encountered  □ unfamiliar | 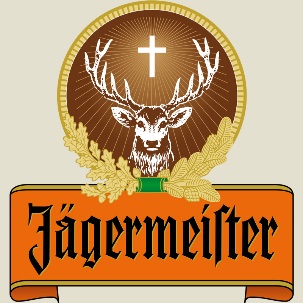 | 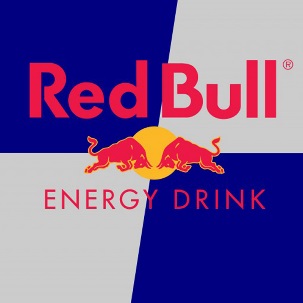 | □ familiar  □ previously encountered  □ unfamiliar |
| □ familiar  □ previously encountered  □ unfamiliar | 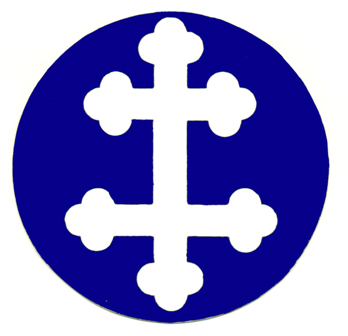 |  |  |

**Supplementary Figure 2** Pre-study survey on the familiarity of the original logos. 15 students, other than the participants included in the study, had to judge 25 original logos as “familiar”, “unfamiliar”, or “previously encountered”.

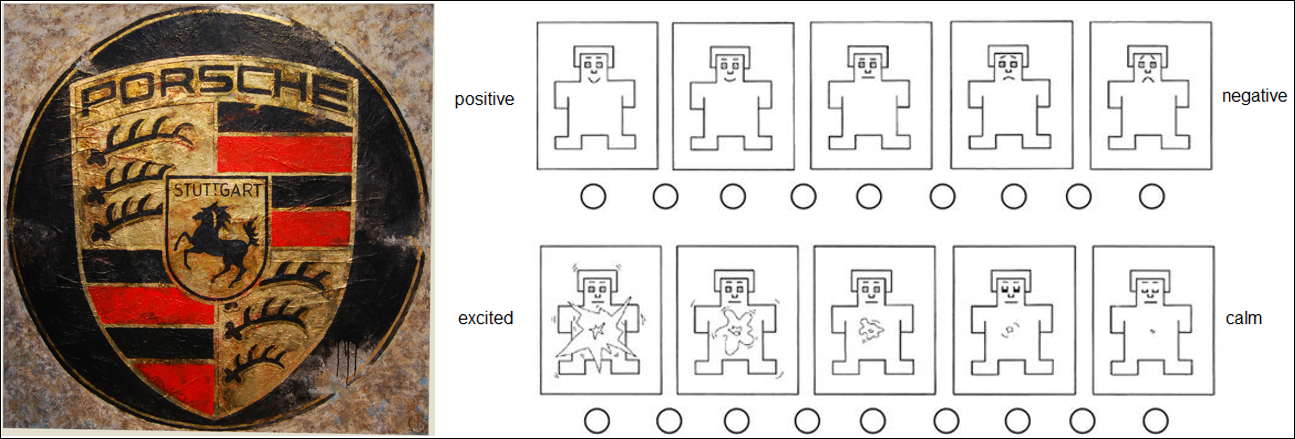

**Supplementary Figure 3** Extract from the self-assessment manikin (SAM). Left side: artistically changed logo of the Porsche brand. Right side: SAM ratings; top row: five pictograms for valence, bottom row: five pictograms for arousal.

| **Subject ___** |  | 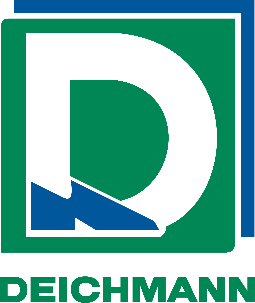 | 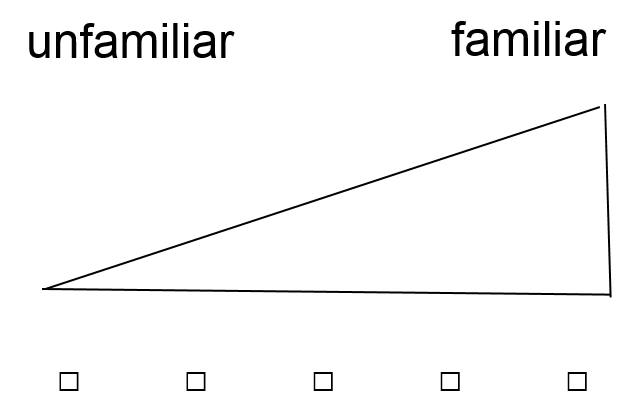 |
| --- | --- | --- | --- |
| 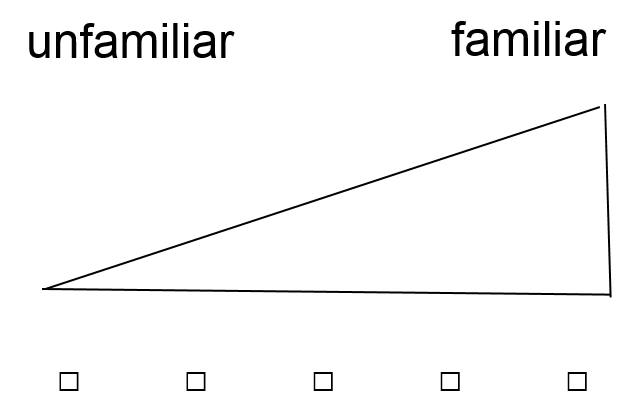 | 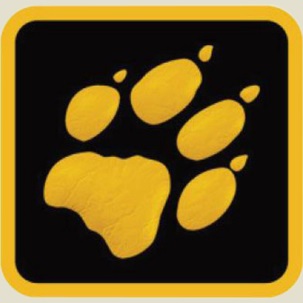 | 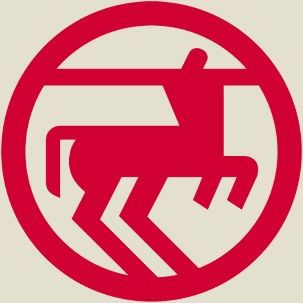 | 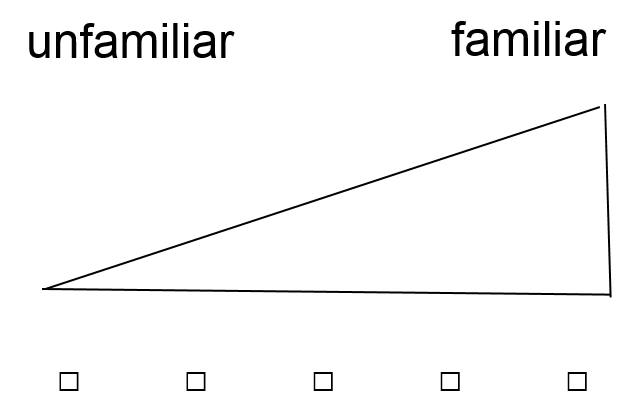 |
| 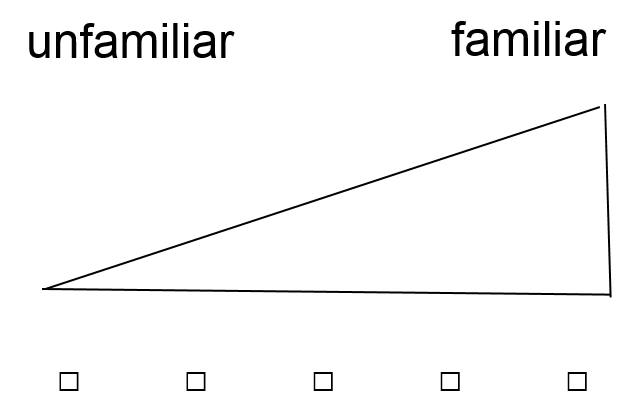 | 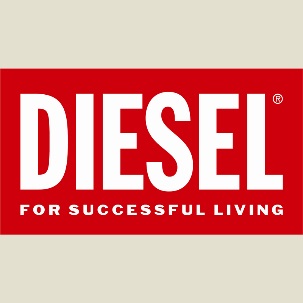 | 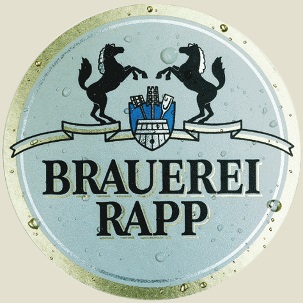 | 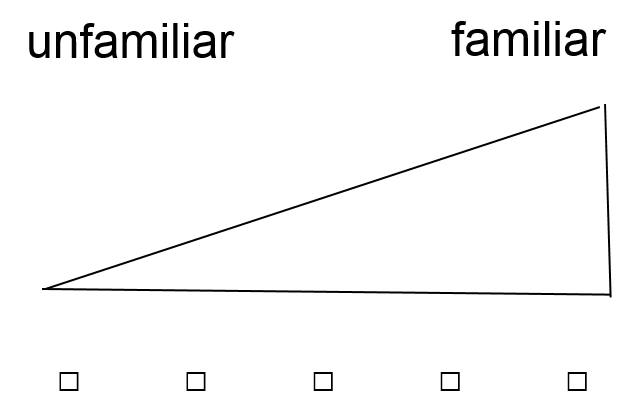 |
| 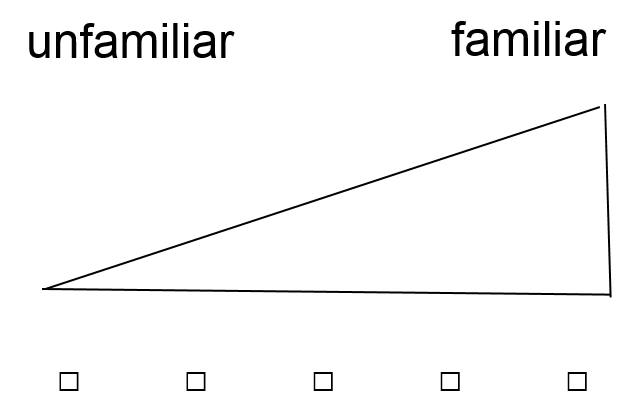 | 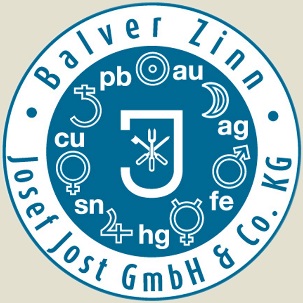 | 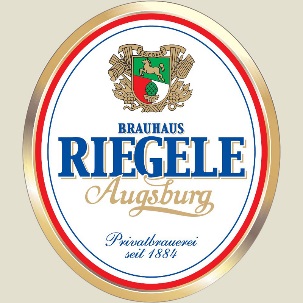 | 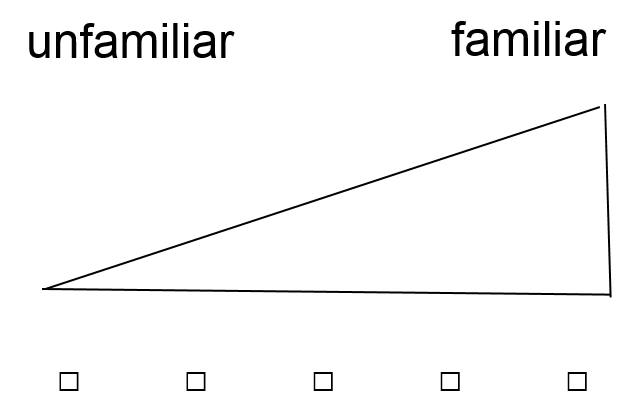 |
| 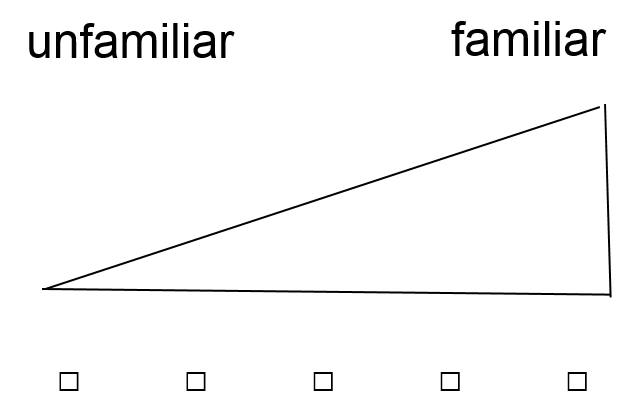 | 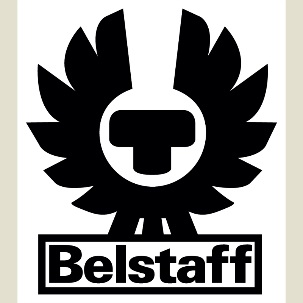 | 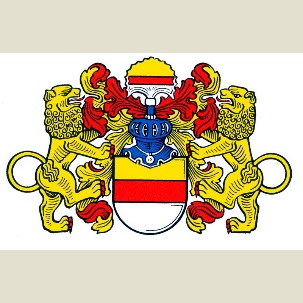 | 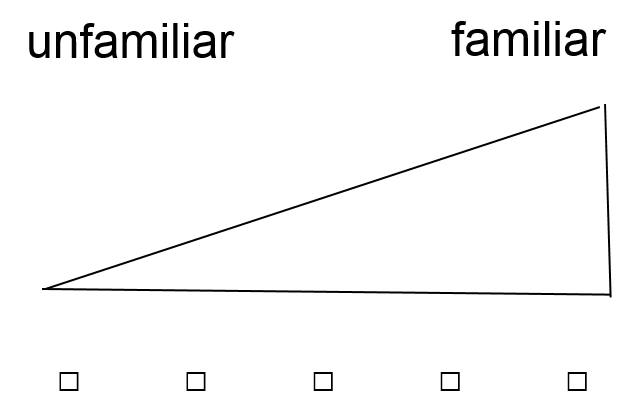 |
| 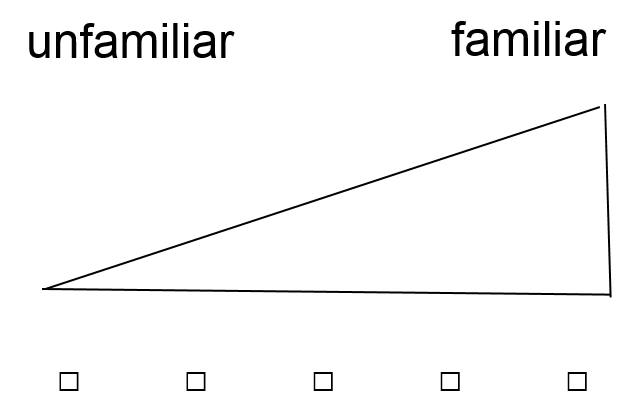 | 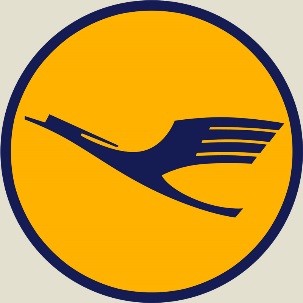 | 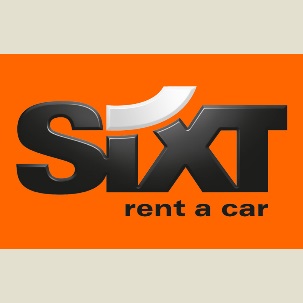 | 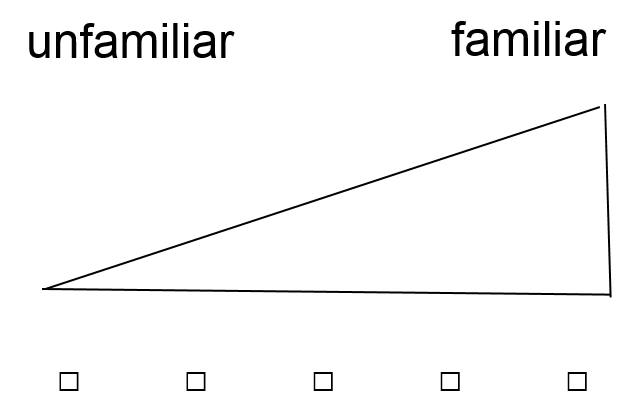 |
| 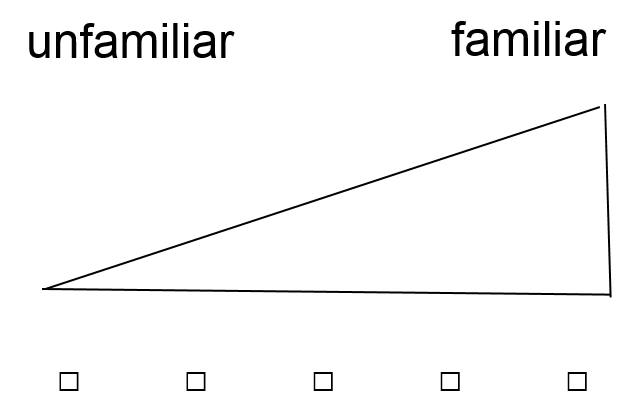 | 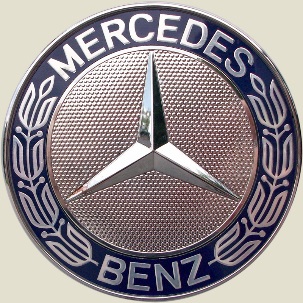 | 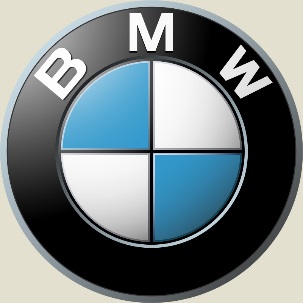 | 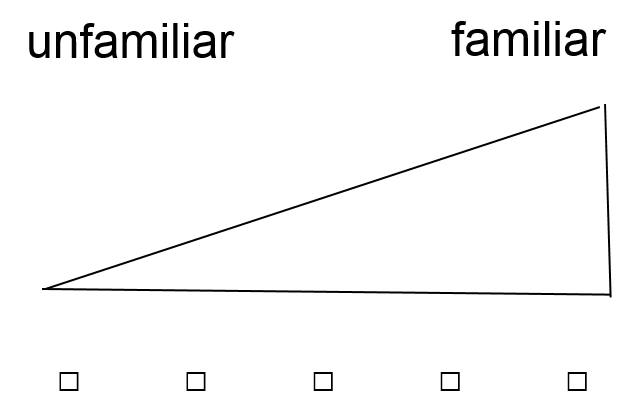 |
| 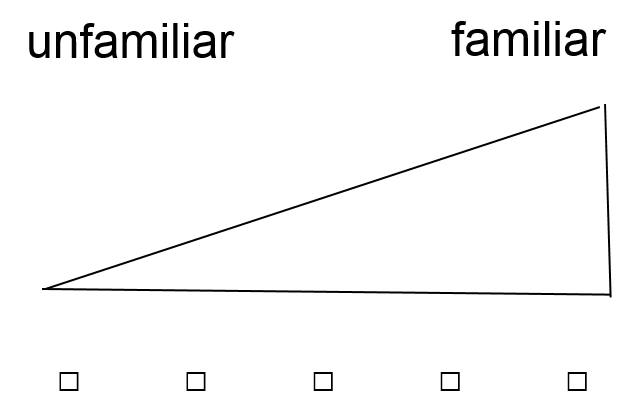 | 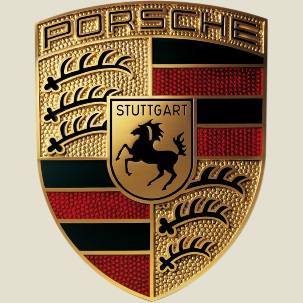 | 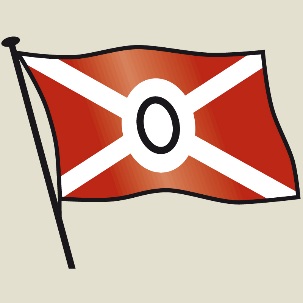 | 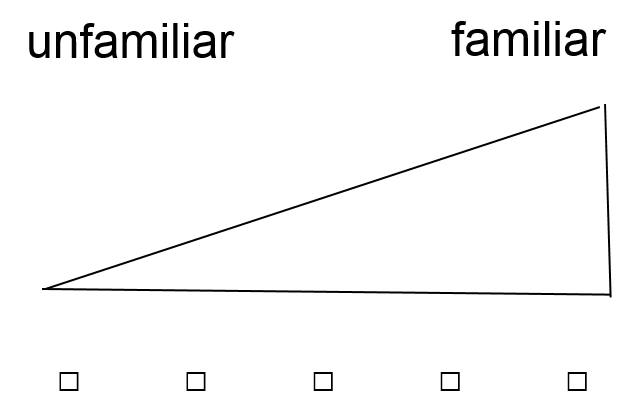 |
| 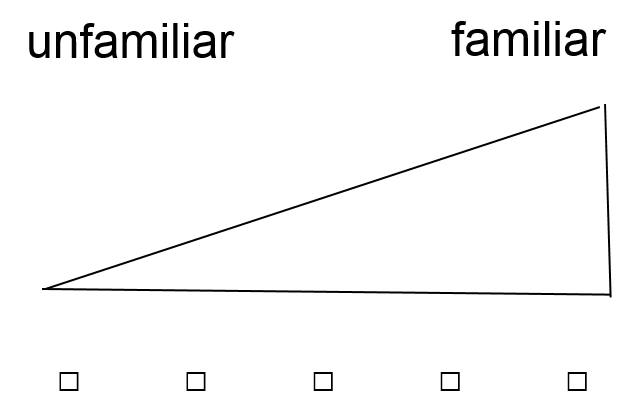 | 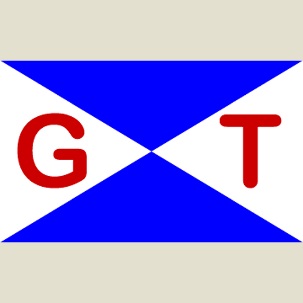 | 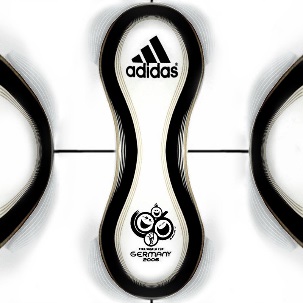 | 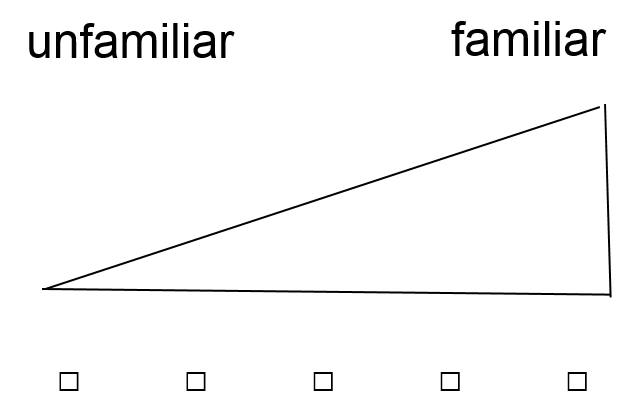 |
| 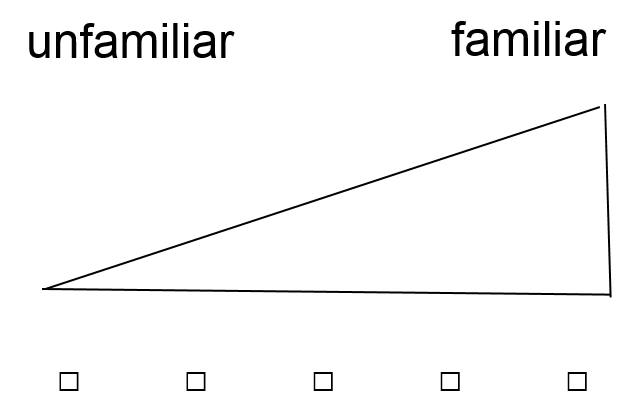 | 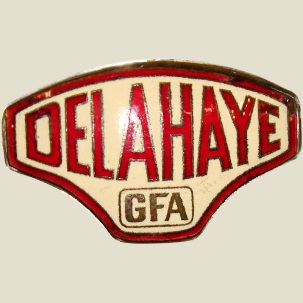 | 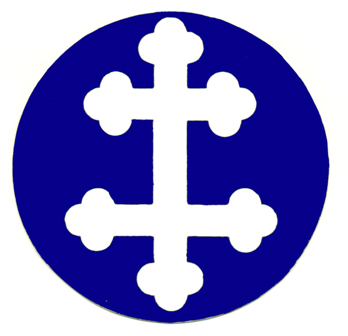 | 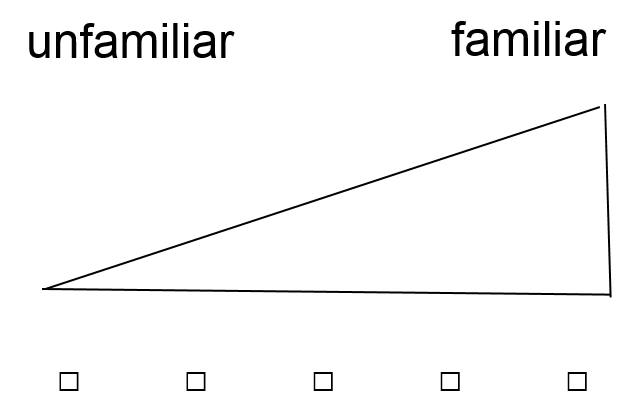 |
| 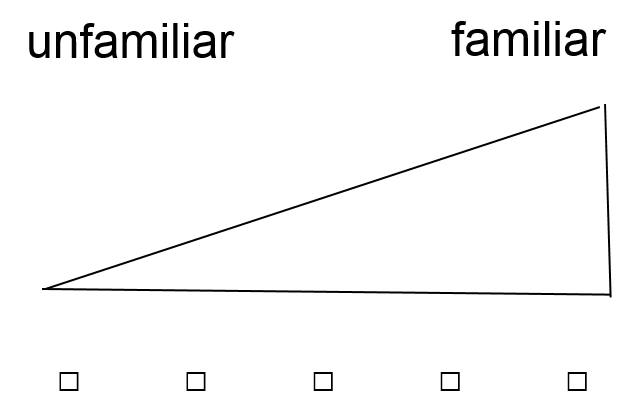 | 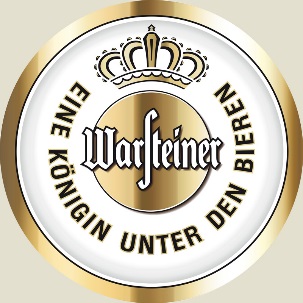 | 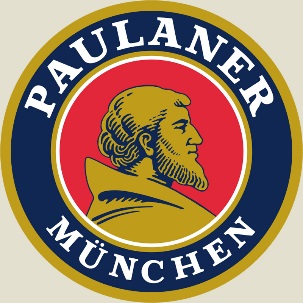 | 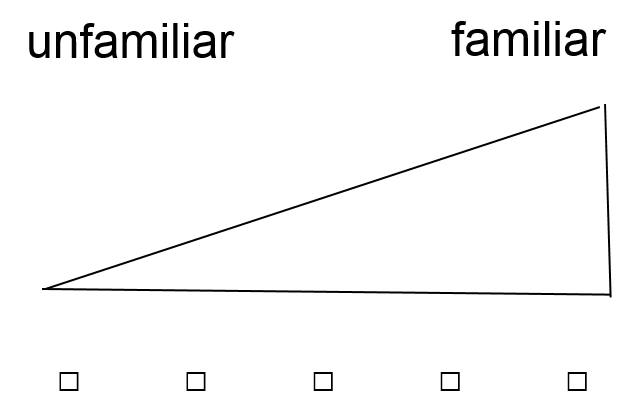 |
| 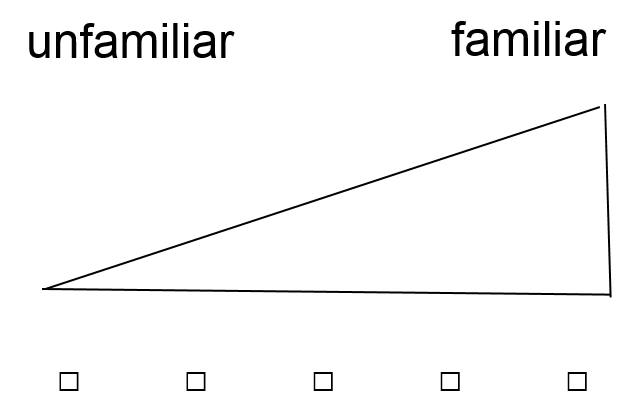 | 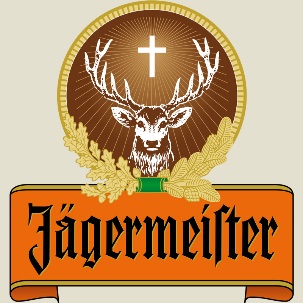 | 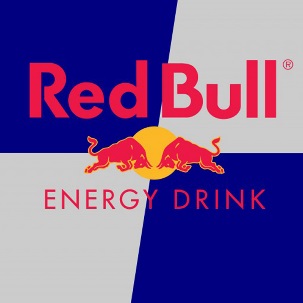 | 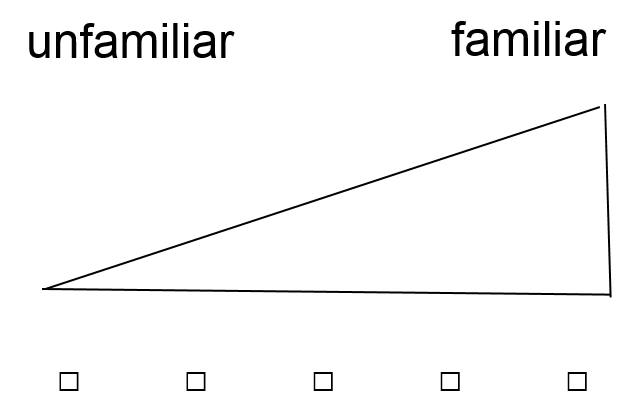 |
| 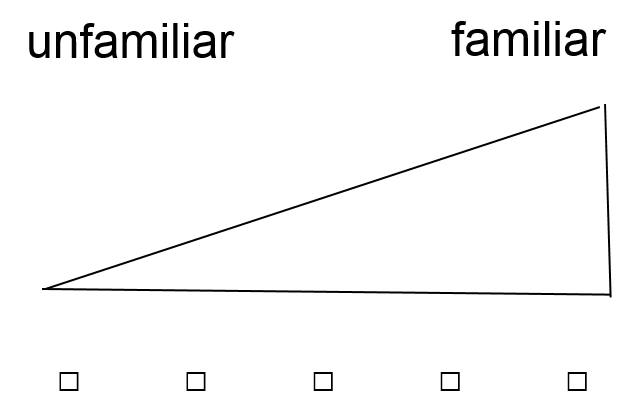 | 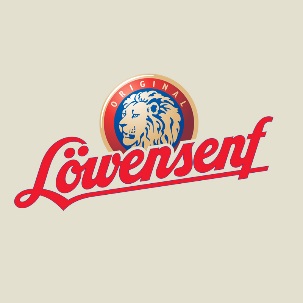 | 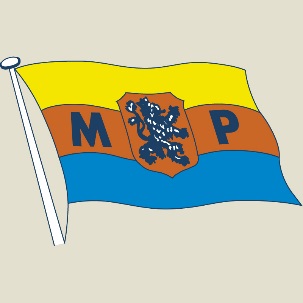 | 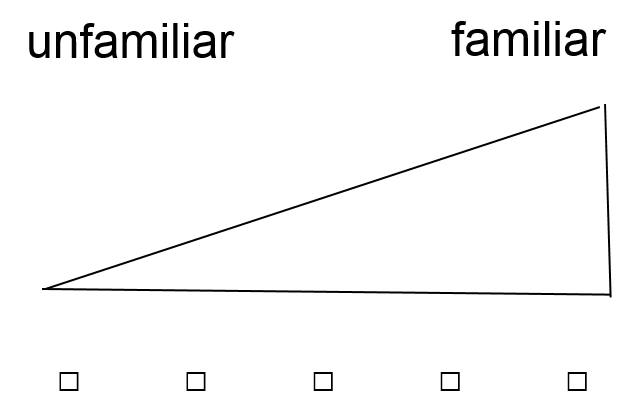 |

**Supplementary Figure 4** Familiarity judgement of original logos on a five-point Likert scale in graphic form.

|  | Familiarity of the original logos |
| --- | --- |
| Jack Wolfskin | 5 |
| Diesel | 4.73333 |
| Balver Zinn | 1.33333 |
| Belstaff | 1.33333 |
| Lufthansa | 4.93333 |
| Deichmann | 4.93333 |
| Rossmann | 4.26667 |
| Brauerei Rapp | 1.66667 |
| Riegele | 2 |
| Münsterwappen | 2.46667 |
| Sixt | 4.8 |
| Mercedes | 4.73333 |
| Porsche | 4.86667 |
| German Tanker | 1.4 |
| Delhaye | 1.4 |
| Warsteiner | 4.86667 |
| Jägermeister | 4.93333 |
| Löwensenf | 4.8 |
| BMW | 5 |
| Orion Bulkers | 1.33333 |
| Adidas | 4.53333 |
| Lorraine Dietrich | 1.46667 |
| Paulaner | 4.86667 |
| Red Bull | 4.93333 |
| Meyer Werft | 1.4 |

Supplementary Figure 5 Evaluation of the familiarity of the original logos on a scale from 1–5. The mean values and standard deviations are shown. Logos which were rated as “familiar” are colored in blue, those rated as “unfamiliar” in orange. Values > 3 indicate a familiar logo and values < 3 an unfamiliar logo.
